# *Leishmania mexicana Centrin* Knock out Parasites Promote M1-polarizing Metabolic Changes

**DOI:** 10.1101/2022.09.16.508215

**Authors:** Greta Volpedo, Timur Oljuskin, Nazli Azodi, Shinjiro Hamano, Greg Matlashewski, Sreenivas Gannavaram, Hira L. Nakhasi, Abhay R. Satoskar

## Abstract

Leishmaniasis is a tropical disease present in more than 90 countries. Presently, there is no approved vaccine for human use. We have previously developed live attenuated *L. mexicana Cen^−/−^ (LmexCen^−/−^)* as a vaccine candidate that showed excellent efficacy that was characterized by reduced activation of Th2 responses and enhanced Th1 responses, contrary to wild type *L. mexicana (Lmex*WT) infection. Towards understanding the interplay between immune mechanisms of protection and metabolic reprogramming, we applied untargeted mass spectrometric analysis to *LmexCen^−/−^* and compared them with *Lmex*WT infection. Data showed that enriched pentose phosphate pathway (PPP) in ears immunized with *LmexCen^−/−^* parasites, compared to naïve and *Lmex*WT-infected ears. This pathway is known to promote an M1 phenotype in macrophages, suggesting a switch to a pro-inflammatory phenotype following *LmexCen^−/−^* inoculation. Accordingly, inhibition of the PPP in macrophages cultured with *LmexCen^−/−^* parasites led to diminished production of nitric oxide, IL-12, and IL-1β, hallmarks of classical activation. Overall, our study revealed novel immune regulatory mechanisms that may be critical for the induction of protective immunity.

## INTRODUCTION

Leishmaniasis is a protozoan disease with a prevalence of 12 million individuals in more than 90 countries worldwide^1–3^. The most common form is cutaneous leishmaniasis (CL), characterized by skin lesions that can ulcerate and give rise to disfiguring scars, which often attract social stigma^1,2,4^. *Leishmania (L*.*) mexicana* is the most prevalent causative agent of cutaneous leishmaniasis (CL) in the New World^5,6^. Despite the elevated morbidity of CL, there are currently no approved human vaccines and the current therapeutic strategies often present significant side effects^7–9^. Our group has previously developed genetically modified *centrin*-deficient *Leishmania* parasites that showed excellent safety and efficacy characteristics in pre-clinical studies^10,11^. *L. major centrin^−/−^* is now under GMP development, however it is unclear whether this Old World strain will be approved as a vaccination strategy or provide cross protection against American *Leishmania* species, as *L. major* is not endemic in the New World. For these reasons we have developed *L. mexicana* mutants (*LmexCen^−/−^*) using the CRISPR/Cas9 technique as a candidate vaccine for American leishmaniasis^12^. *LmexCen^−/−^* parasites presented a growth defect, which resulted in loss of virulence *in vitro* and *in vivo*. Furthermore, immunization with *LmexCen^−/−^* did not cause cutaneous lesions in susceptible mouse models and was efficacious against challenge with the virulent strain by eliciting strong protective immunity^12^.

As we continue investigating *LmexCen^−/−^* parasites as a vaccination strategy, we are interested in understanding the immune determinants of protection and in particular the early drivers of vaccine-mediated immunological profiles. It is well established that *L. mexicana* infection can prevent the induction of a pro-inflammatory response by disrupting IL-12 production^13^ and Th1 differentiation^14^. Furthermore, *L. mexicana* is known to induce expression of arginase and polarize macrophages towards an M2 anti-inflammatory phenotype^15^. On the other hand, *LmexCen^−/−^* parasites can induce IL-12 production in macrophages, important to polarize the host cell towards an M1 pro-inflammatory phenotype and induce a Th1 response^12^. While M2 macrophages are usually associated with susceptibility to CL, M1 macrophages can mediate protection via their antimicrobial functions^16^. Early differentiation into M1 or M2 macrophages can have a profound impact on the development of an effective adaptive immune response, however, the molecular mechanisms by which *LmexCen^−/−^* parasites lead to classical activation of macrophages have not yet been explored.

A growing body of evidence shows that metabolic changes in macrophages infected with *Leishmania* parasites can impact the availability of nutrients necessary for amastigote replication and affect the cell’s polarization and host potential^17^. For instance, M1 macrophages in *Leishmania* dermal granulomas upregulate the pentose phosphate pathway (PPP), important for reactive oxygen species (ROS) production^17^ and M1 effector responses^18^. The PPP has also been shown to control the replication of *Trypanosoma cruzi*, another protozoan parasites, in macrophages by promoting ROS and nitric oxide (NO) release^19^. Interestingly, upregulation of the PPP in macrophages occurs within one hour of lipopolysaccharide (LPS) stimulation, and several hours prior to the induction of pro-inflammatory mediators such as superoxide, IL-1β, TNF-α, IL-6, and IL-12^20^. These observations highlight the temporal pattern of macrophage activation and suggest that metabolic changes occur prior to, and may cause, immunological changes.

Recent studies have started exploring the role of metabolic changes as drivers of immunomodulation in the context of vaccination. For instance, neutrophils, monocytes, and macrophages undergo metabolic reprogramming following immunization with Bacillus Calmette-Guérin (BCG), a live attenuated vaccine for tuberculosis, which results in a protective immune response^21,22^. In macrophages, this immune response is specifically induced by the activation of aerobic glycolysis^22^. These studies suggest an important role for metabolic reprogramming in vaccination-mediated immunity against pathogens.

Interestingly, studies with other pathogens show that vaccinated individuals display defined metabolic signatures compared to those who did not receive the vaccine, which could be used as reliable biomarkers of protection and vaccine efficacy^23^. Live attenuated vaccines have also been shown to harbor significant metabolic differences compared to their parental strains, as in the case of *Mycobacterium tuberculosis* and the live attenuated MTBVAC^24^. Different metabolic profiles in parental and mutant strains could translate into metabolic changes within the host cell depending on exposure to the vaccine or the virulent pathogen. These observations highlight how determining pathogen and strain-specific metabolic reprogramming, as well as their subsequent effects on biological and immunological downstream processes, is a useful tool to identify and characterize unique biomarkers of vaccination.

Towards the goal of uncovering specific metabolic signatures associated with vaccination, our group has sought to explore the immune determinants underlying protection after immunization with *Cen^−/−^ Leishmania* mutants. In particular, we previously showed that immunization with *LmexCen^−/−^* leads to enhanced Th1 responses in C57BL/6 mice^12^. In this study, we aim to investigate some of the metabolic drivers of these immunological changes. We explored the metabolic profiles occurring locally at the inoculation site after immunization with *LmexCen^−/−^* to understand where some of the immunological responses originate, which we have previously reported. We were also interested in comparing *LmexCen^−/−^* immunization with *LmexWT* infection to investigate whether there are any differences in the metabolic changes that can explain the divergent immune responses we observe in vaccinated compared to control mice. We chose a 7-day time point to specifically explore the early changes that can shape the immune responses and drive resistance or susceptibility. We found several metabolic pathways enriched in *LmexCen^−/−^* infected ears that are consistent with the protective immune response observed after immunization. To our knowledge, this is the first report utilizing metabolomics to investigate metabolomic drivers of immune responses in the context of a *Leishmania* vaccine.

## RESULTS

### Ear tissue of mice immunized with *LmexCen^−/−^* displays metabolic differences compared to naïve mice and mice infected with *Lmex*WT

We collected the ear tissue from mice immunized with *LmexCen^−/−^*, infected with *Lmex*WT, and naïve mice and performed untargeted mass spectrometry analysis to identify unique metabolic signatures of disease and vaccination (Figure 1). Using volcano plots, which plot fold change in the x axis and significance in the y axis, we selected significant features from the positive (Figure 2a) and negative (Figure 2b) modes of mass spectrometry analysis. The peaks identified with the positive mode correspond to protonated molecules, while those identified with the negative mode correspond to deprotonated analytes. A fold change threshold (x) 2 and t-tests threshold (y) 0.05 were set for the volcano plots (Figure 2a-b). We have then used significant features from the volcano plots for further analysis. We have also used partial least squares-discriminant analysis (PLS-DA), a supervised multivariate analytic approach, to select metabolites unique to our three conditions. As shown in Figure 2 c and d for the positive and negative modes, respectively, naïve mice and the groups inoculated with either *LmexCen^−/−^* or *Lmex*WT each displayed unique metabolic signatures, highlighting a significant *Leishmania*-specific metabolic shift with distinct characteristics depending on whether the mice were exposed to the mutant or parental strain.

**Figure 1.**
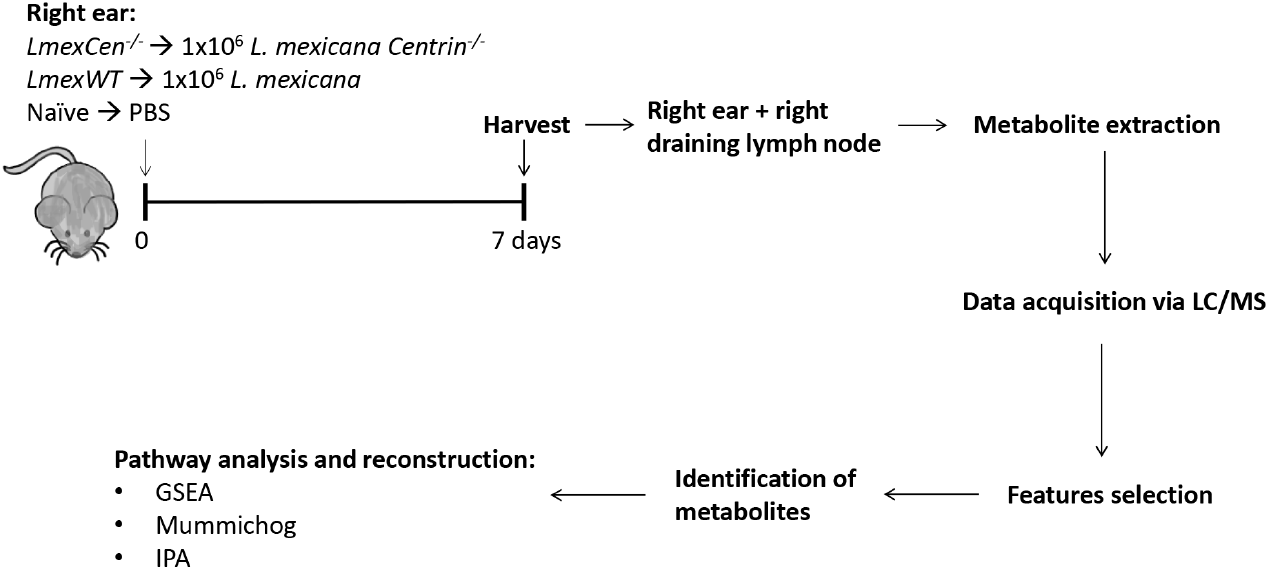
Experimental design for metabolomic analysis of infection with *Lmex*WT and immunization with *LmexCen^−/−^*. Female C57BL/6 mice were injected intradermally in the ear with stationary phase *Lmex*WT, *LmexCen^−/−^* parasites, or PBS. After 7 days, the injected ears and their respective draining lymph nodes were collected and processed for mass spectrometry. The data acquired was used to identify enriched metabolites and determine activated pathways in the tissue. N=5 for each group. Abbreviations: LC/MS: liquid chromatography / mass spectrometry; GSEA: gene set enrichment analysis; IPA: ingenuity pathway analysis.

**Figure 2.**
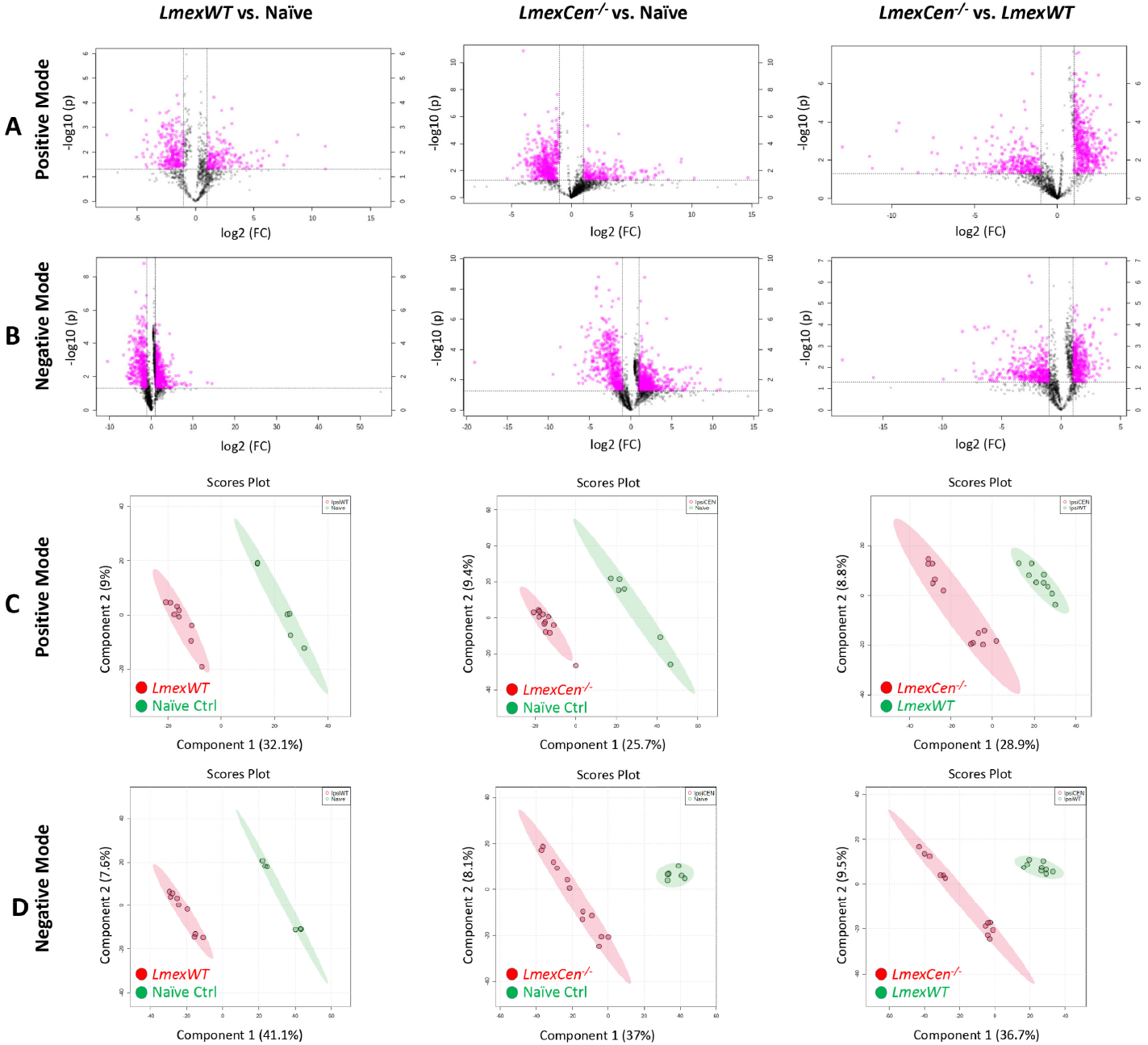
Infection with *Lmex*WT and immunization with *LmexCen^−/−^* display different metabolic signatures in at the inoculation site. Normalized data from ear tissue from C57BL/6 mice after 7 days of infection with *Lmex*WT, immunization with *LmexCen^−/−^*, or naïve control was used to perform statistical analysis. **A, B)** Features selected by volcano plot from positive (**A**) and negative (**B**) modes for ear tissue of *Lmex*WT vs. naïve, *LmexCen^−/−^* vs. naïve, and *LmexCen^−/−^* vs. *Lmex*WT mice using LC/MS with fold change threshold (x) 2 and t-tests threshold (y) 0.05. Both fold changes and p-values are log transformed. **C, D)** Partial least squares-discriminant analysis (PLS-DA) from positive (**C**) and negative (**D**) mode for ear tissue of *Lmex*WT vs. naïve, *LmexCen^−/−^* vs. naïve, and *LmexCen^−/−^* vs. *Lmex*WT mice.

### Immunization with *LmexCen^−/−^* leads to M1-polarizing metabolic changes at the inoculation site

In order to identify specific metabolic pathways enriched in mice immunized with *LmexCen^−/−^*, we used both mummichog and gene set enrichment analysis (GSEA) analysis. Mummichog analysis uses an algorithm that predicts functional activity bypassing metabolite identification, while GSEA requires the metabolites to be identified before pathway/network analysis. We used the Integrated MS Peaks to Pathways plot to summarize the results of the Fisher’s method for combining mummichog (y) and GSEA (x) p-values from the positive (Figure 3 a) and negative (Figure 3 b) mode data sets, indicating the metabolic pathways enriched in the ear tissue of different groups. In particular, we were interested in identifying any significantly enriched pathways known to promote a pro-inflammatory protective immune response in immunized compared to naïve mice. Based on our analysis, we identified aspartate metabolism and pentose phosphate pathway (PPP), which are involved in M1-polarization of macrophages ^17,20,25^. In particular, our results showed enriched alanine, aspartate, and glutamate metabolism in the ears of *LmexCen^−/−^* immunized mice, compared to naïve mice as highlighted by the black box in Figure 3.3 a. Furthermore, the PPP was also enriched in the ears of *LmexCen^−/−^* immunized mice, compared to naïve and *Lmex*WT-infected mice (Figure 3.3 a-b). Based on the t scores for the matched metabolites, both aspartate metabolism and the PPP were upregulated in the *LmexCen^−/−^* immunized group, revealing a mechanism for protection mediated by *LmexCen^−/−^* parasites. In particular, D-erythrose 4-phosphate, sn-glycerol 3-phosphate, 3-ketolactose, D-glucose, melibiose, and thiamin diphosphate were significantly up-regulated in the infected group, as identified using Metscape (Figure 4). Table 1 shows all PPP mediators enriched in the *LmexCen^−/−^* dataset compared to the naïve dataset.

**Table 1.**
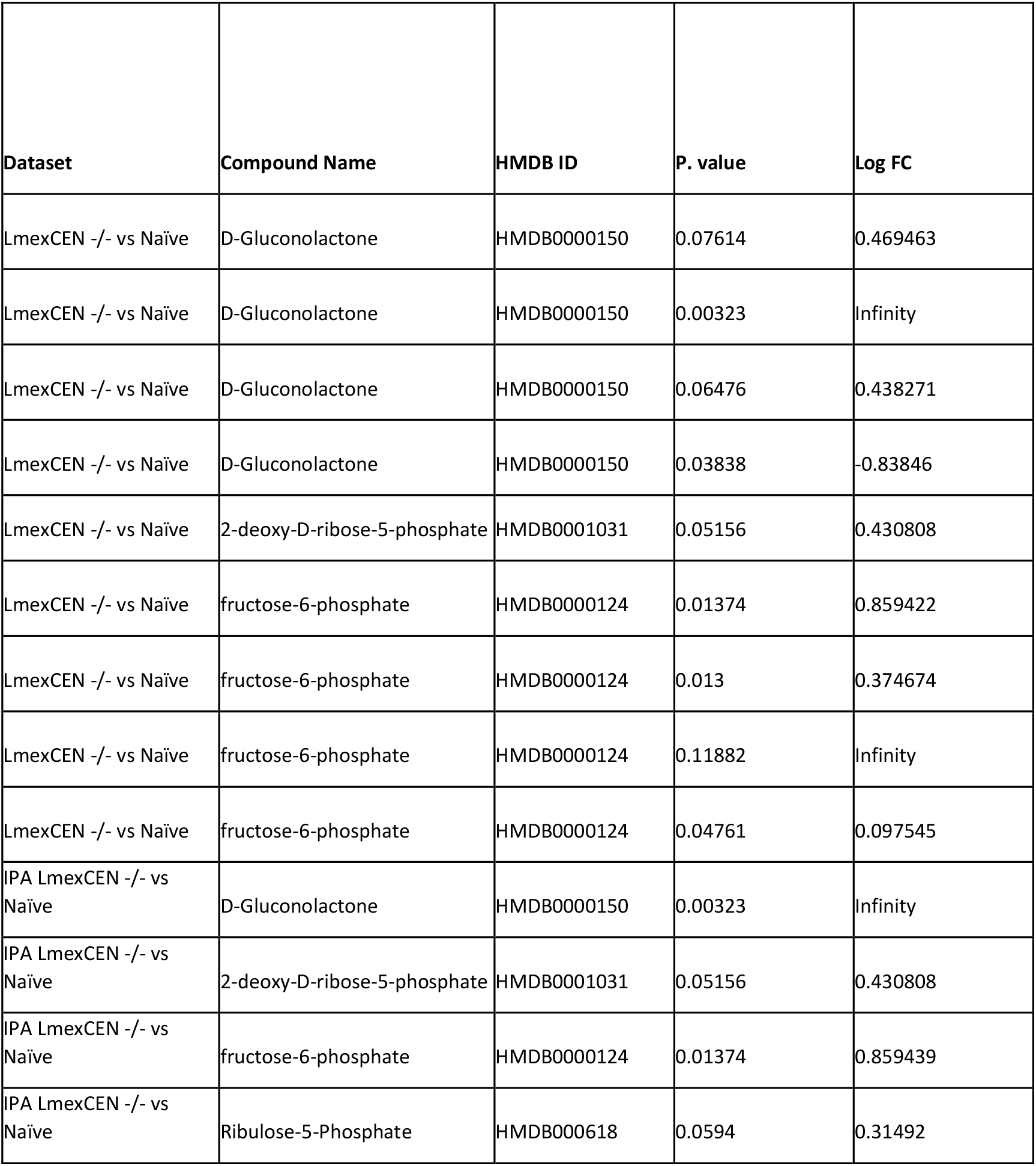
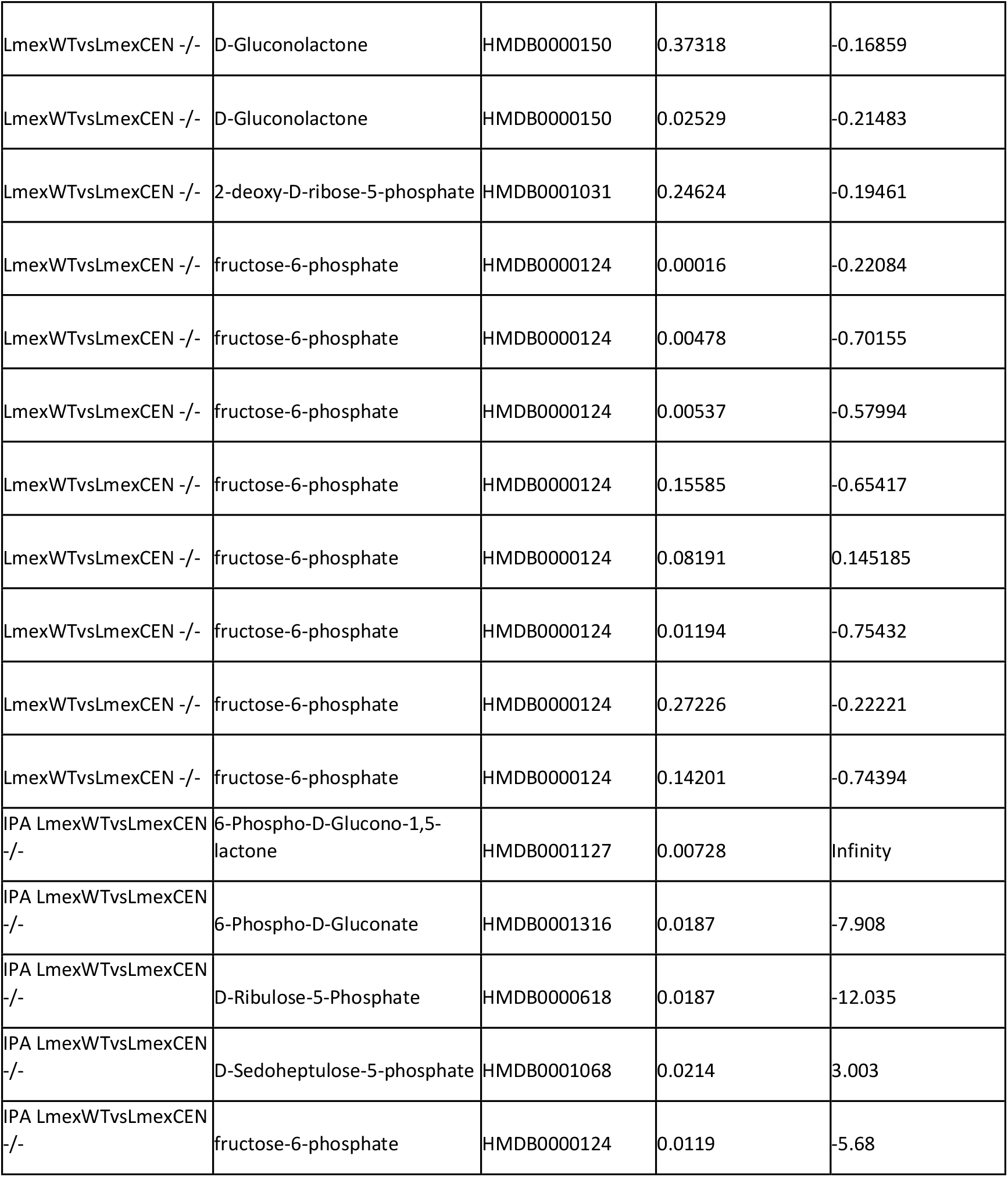
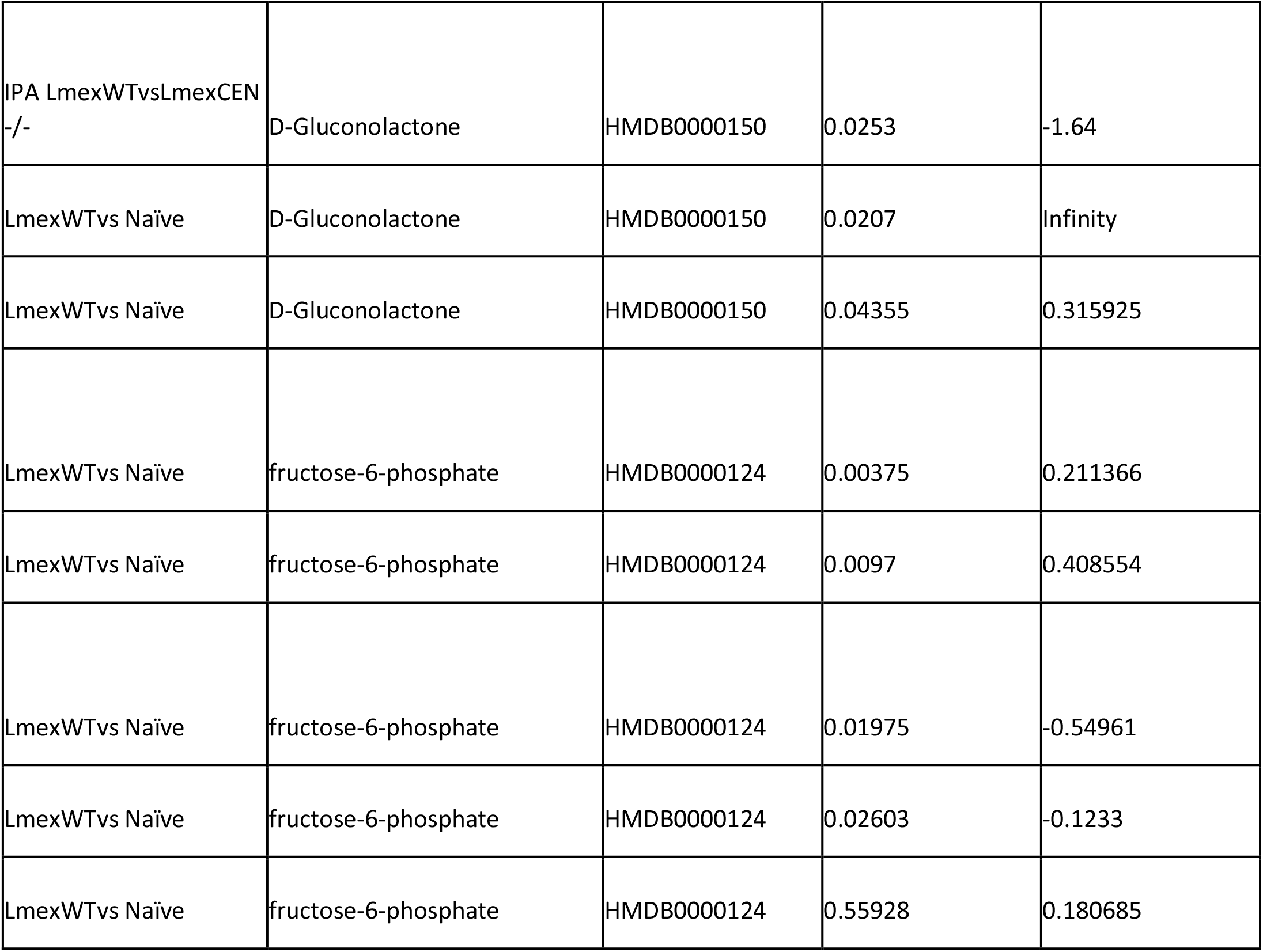
Pentose phosphate pathway mediators enriched in the *LmexCen-/-* and *Lmex*WT datasets. Normalized data ear tissue from C57BL/6 mice inoculated with *LmexCen-/-, Lmex*WT or naïve controls was used to identify pentose phosphate pathway metabolites.

**Figure 3.**
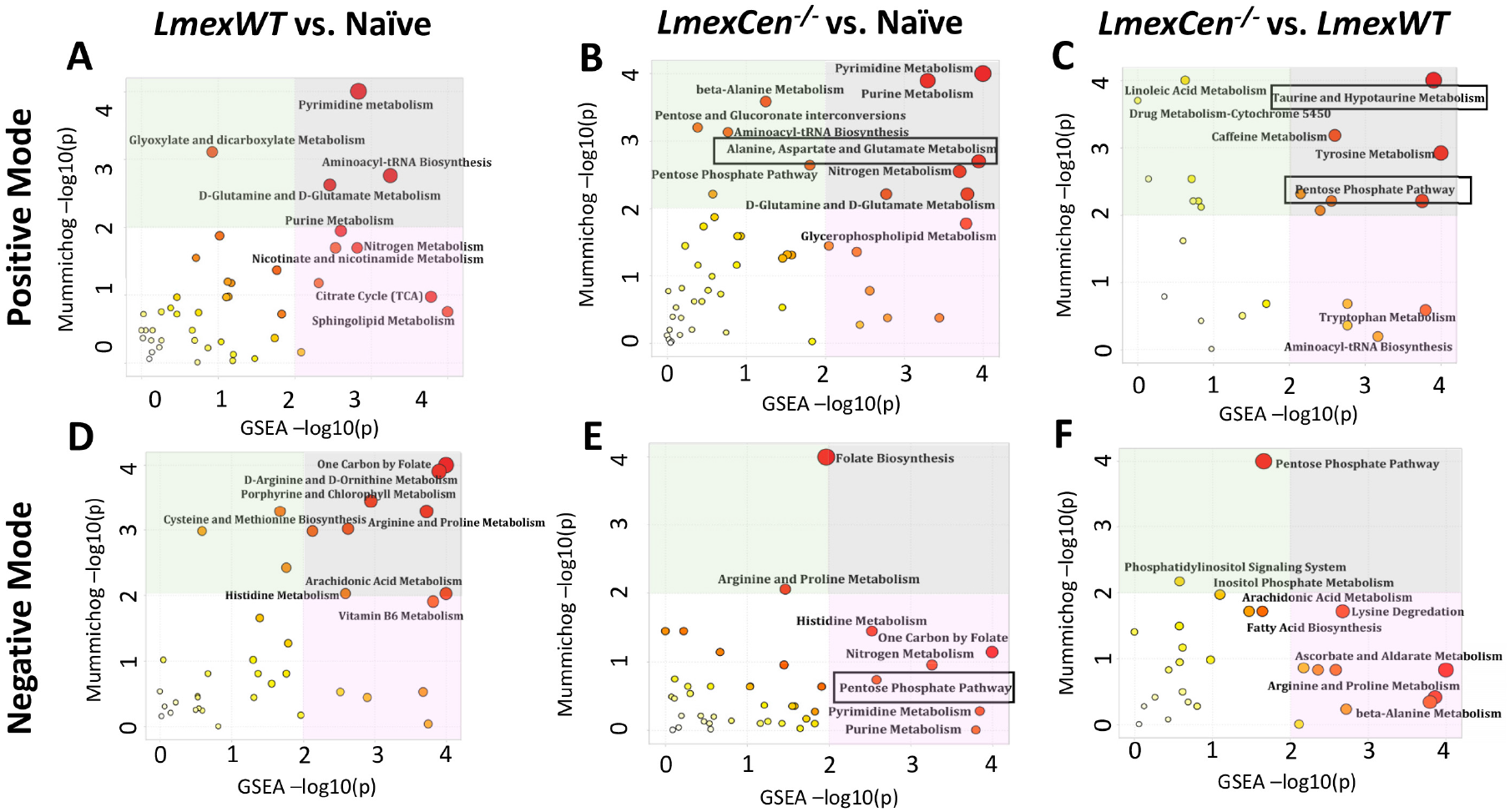
Metabolic pathways enriched in mice infected with *Lmex*WT or immunized with *LmexCen^−/−^* at the inoculation site. Normalized data from ear tissue from C57BL/6 mice after 7 days of infection with *Lmex*WT, immunization with *LmexCen-/-*, or naïve control was used to perform peaks to pathway analysis. Using the MS Peaks to Paths module in MetaboAnalyst5.0, the mummichog and gene set enrichment analysis (GSEA) p-values were combined. The Integrated MS Peaks to Paths plot summarizes the results of the Fisher’s method for combining mummichog (y) and GSEA (x) p-values from the positive (**A**) negative (**B**) mode data sets, indicating the metabolic pathways enriched. The size and color of the circles correspond to their transformed combined p-values. Large and red circles are considered the most perturbed pathways. The colored areas show the significant pathways based on either mummichog (blue) or GSEA (pink), and the purple area highlights significant pathways identified by both algorithms. Highlighted by the black boxes are pathways associated with a protective immune response against leishmaniasis.

**Figure 4.**
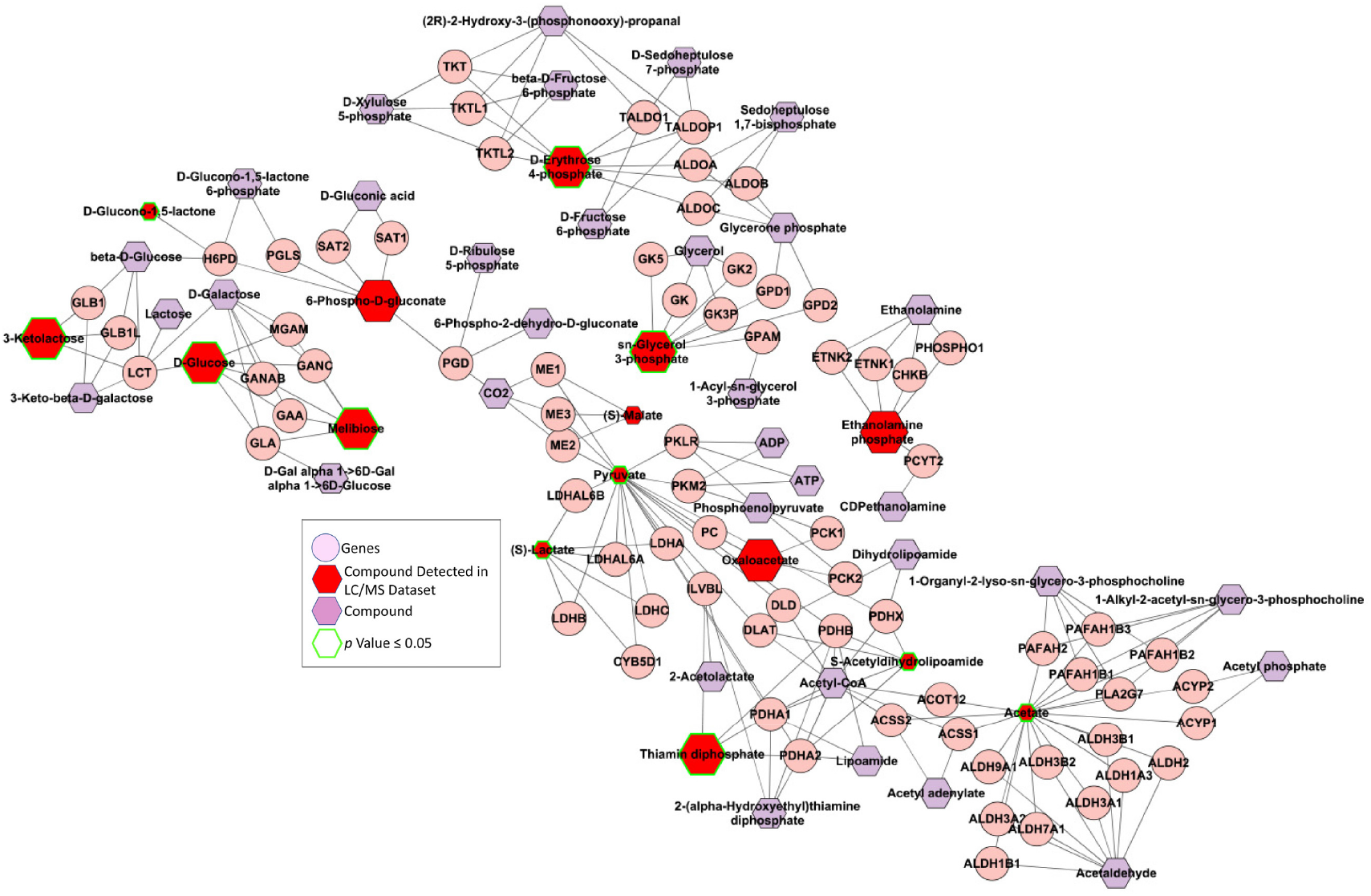
Immunization with *LmexCen^−/−^* leads to enriched Pentose Phosphate Pathway metabolism. Normalized data from ear tissue from C57BL/6 mice after 7 days of infection with *Lmex*WT, immunization with *LmexCen-/-*, or naïve control was used to perform statistical analysis. Metabolite network identified with Metscape. Metabolites of the Pentose Phosphate Pathway are represented in this graph. Larger hexagons represent up-regulation, while smaller hexagons represent down-regulation. Red hexagons represent compounds detected in the data set, while hexagons with a green outline represent statistically significant metabolites (p-value ≤ 0.05). The purple hexagons represent compounds that are associated with the pathway but are not detected in the input dataset. The pink circles represent the genes regulating the biosynthetic activities.

### Co-culture with *LmexCen^−/−^* parasites leads to pentose phosphate pathway-dependent M1 phenotype in macrophages

Our mass spectrometry analysis revealed upregulated PPP in the *LmexCen^−/−^* group, which is known to polarize macrophages towards an M1 phenotype. We have previously observed enhanced pro-inflammatory responses following *LmexCen^−/−^* immunization *in vitro* and *in vivo*^12^, however it is not yet known whether the PPP significantly contributes to driving this immune profile. In order to investigate this, we measured nitric oxide (NO), IL-12, and IL-1β in bone marrow-derived macrophages (BMDMs) cultured with *LmexCen^−/−^* parasites and stimulated with LPS in presence or absence of the PPP inhibitors 6-aminonicotinamide (6-AN) and dehydroepiandrosterone (DHEA) (Figure 5a). NO, IL-12, and IL-1β are hallmark mediators of M1 macrophages induced by the PPP and involved in parasite killing and the induction of Th1 immune responses^16,18,26,27^. Our results show that LPS stimulation lead to a significantly higher NO production in BMDMs cultured with *LmexCen^−/−^* parasites, compared to the uninfected group (Figure 5b). Treatment with either 6-AN or DHEA resulted in a significant reduction in NO in both the uninfected and *LmexCen^−/−^* groups (Figure 5b), suggesting that inhibition of the PPP causes impaired NO production. Incubation with either 6-AN or DHEA also resulted in a significant reduction in IL-12 production in both the uninfected and *LmexCen^−/−^* groups, compared to their respective controls (Figure 5c). The same trend was observed in the *LmexCen^−/−^* group for the production of IL-1β (Figure 5d). Taken together these results highlight a role for PPP in *LmexCen^−/−^*-mediated M1 polarization and pro-inflammatory effector functions in macrophages (Figure 6).

**Figure 5.**
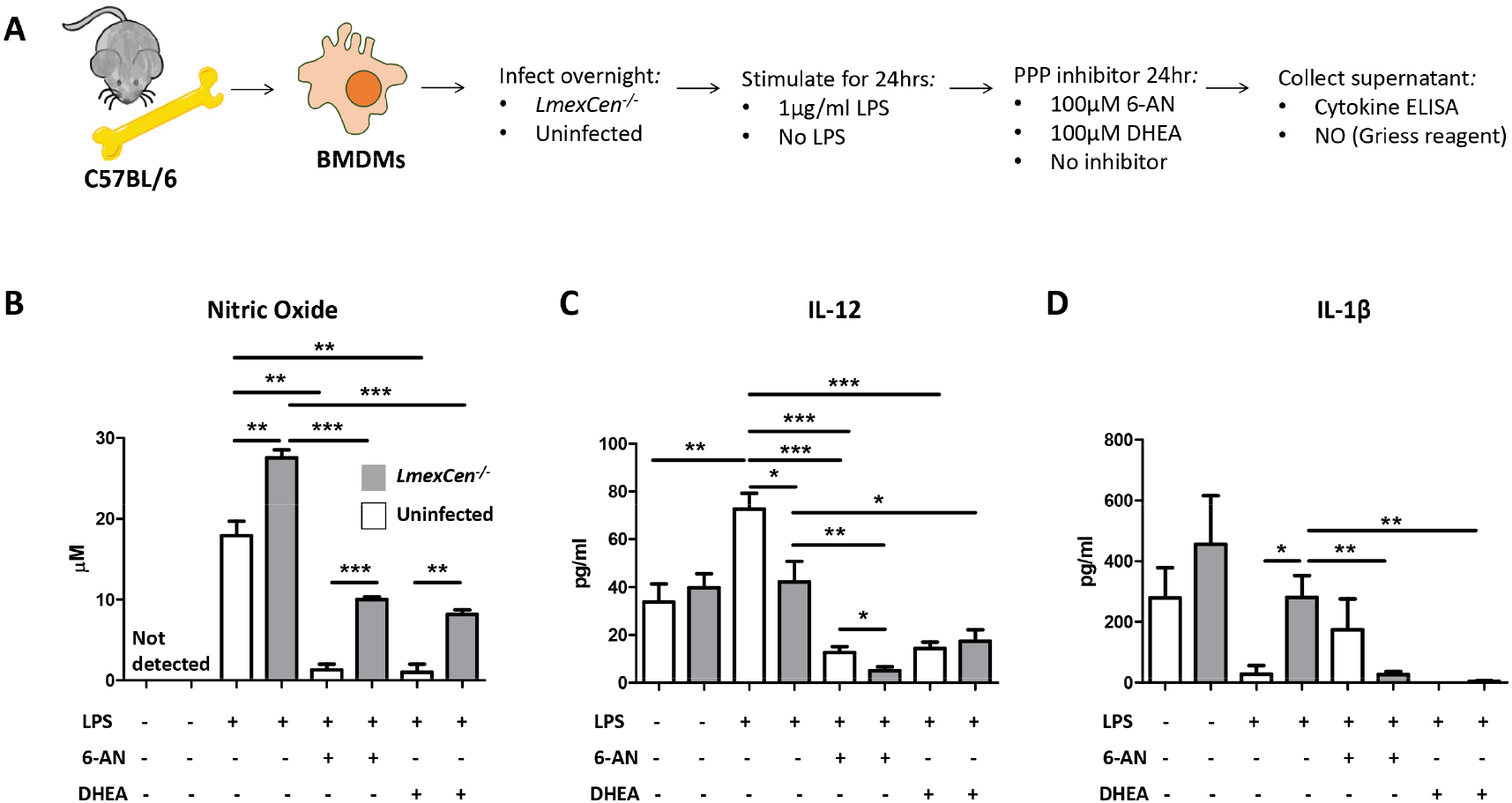
Pentose phosphate pathway-dependent nitric oxide, IL-12, and IL-1β release in BMDMs. **A**) BMDMs were extracted from C57BL/6 mice and cultured with medium or *LmexCen^−/−^* and stimulated *in vitro* by LPS for 24 hrs, and 6-AN or DHEA for 24 hrs. **B**) Nitric oxide production was determined by Griess reaction. IL-12 (**C**) and IL-1β (**D**) production were determined by cytokine ELISA. Data represents one of three experiments with N=3-5 per group. *P < 0.05, ** P ≤ 0.01 and *** P ≤ 0.001 unpaired t test. Error bars represent SEM. Abbreviations: LPS, lipopolysaccharide; 6-AN, 6-aminonicotinamide; DHEA, dehydroepiandrosterone; NO, nitric oxide.

**Figure 6.**
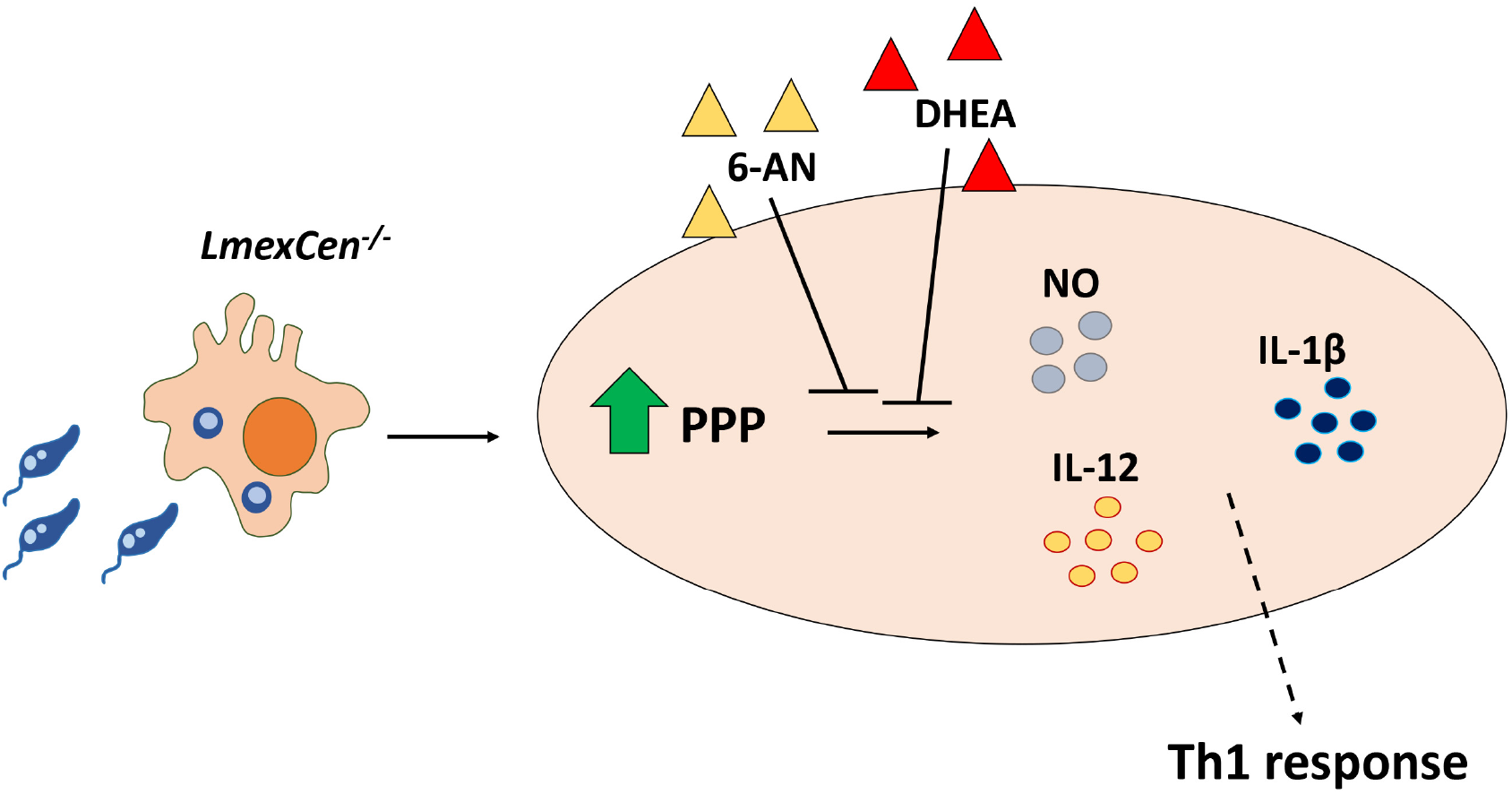
Immunization with *LmexCen^−/−^* parasites leads to induction of pentose phosphate pathway. Graphical scheme of *LmexCen^−/−^*-induction of the PPP, resulting in the production of NO, IL-12, and IL-1b in macrophages. Abbreviations: 6-AN, 6-aminonicotinamide; DHEA, dehydroepiandrosterone; NO, nitric oxide.

### *Leishmania-centrin* deficiency results in differential host metabolic reprogramming, compared to *Lmex*WT infection

After identifying metabolic drivers of immune protection in immunized compared to non-immunized models, we explored the host metabolic differences mediated by *LmexCen^−/−^*, compared to *Lmex*WT parasites. Interestingly, arginase metabolism, and PPP identified in the ears of immunized mice, were not enriched in the *Lmex*WT-infected group compared to naïve mice (Figure 3 a-b), highlighting differential metabolic regulation following exposure to the mutant or parental strain of *L. mexicana*. Table 1 shows all PPP mediators enriched in the *LmexCen^−/−^* dataset compared to the *Lmex*WT dataset.

Our combined mummichog and GSEA analysis also revealed the enrichment of taurine and hypotaurine metabolism in ears immunized with *LmexCen^−/−^*, compared to those infected with *Lmex*WT parasites (Figure 3a). This pathway is known to play a role in the healing of CL lesions^28^ and it was upregulated in the *LmexCen^−/−^*-group, based on the t scores of the matched metabolites. In particular, the taurine/hypotaurine metabolite glutarine was significantly upregulated in *LmexCen^−/−^*-immunized mice, compared to *Lmex*WT-infected mice.

A Venn diagram of the metabolic pathways enriched in the ears of mice infected with *Lmex*WT or immunized with *LmexCen^−/−^*, compared to uninfected naïve mice, can be found in Figure 7. Despite a handful of common pathways, the majority of metabolic changes seems to be strain-specific, suggesting that *centrin* may play a role in this divergent metabolic regulation of the mammalian host cell.

**Figure 7.**
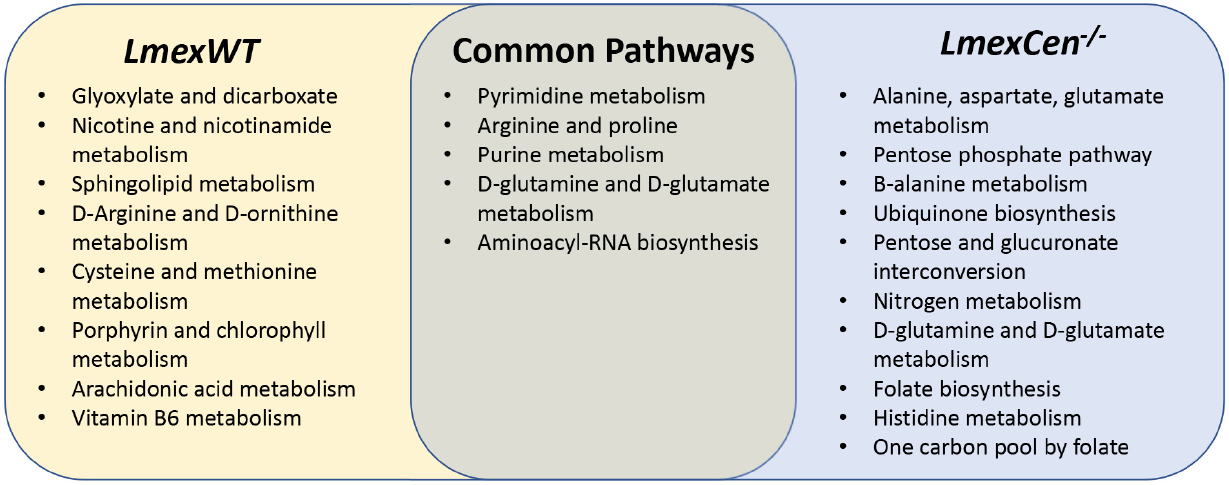
Unique and common pathways enriched in the ears following *LmexCen-/-* immunization or *Lmex*WT infection. Venn diagram of the metabolic pathways significantly enriched based on mummichog and GSEA analysis in the ear of C57BL/6 mice after 7 days of infection with *Lmex*WT or immunization with *LmexCen-/-*, compared to uninfected naïve mice.

Taken together, our results show that *LmexCen^−/−^* parasites lead to differential metabolic reprogramming, compared to the WT strain. In particular, the *LmexCen^−/−^* mutants promote metabolic profiles known to drive protective immunity against leishmaniasis.

### Draining lymph nodes display different metabolic reprogramming, compared to cutaneous tissue, during early exposure to *Leishmania*

After exploring metabolic reprogramming locally at the inoculation site, we were interested in investigating metabolic changes in the draining lymph nodes in the early stages of the infection. Similarly, to the process we used for the ear tissue, volcano plots were employed to select significant features from the positive (Supplementary Figure 1a) and negative (Supplementary Figure 1b) modes, and PLS-DA to identify variations amongst the experimental groups from the positive (Supplementary Figure 1c) and negative (Supplementary Figure 1d) modes. Overall, our results show distinct metabolic signatures in the draining lymph nodes of mice inoculated with either *LmexCen^−/−^* or *Lmex*WT, and naïve controls, in a manner comparable to what we observed locally in the ear skin tissue.

Interestingly, mass spectrometry analysis of lymph node tissue revealed different results from what we found locally in the ears. While we did not observe enrichment of the same M1-promoting pathways in *LmexCen^−/−^*-immunized mice, our data shows enriched linoleic acid metabolism in the group immunized with *LmexCen^−/−^* mutants, compared to naïve mice (Supplementary Figure 2a-b). Based on the t scores for the matched metabolites, this pathway was activated in the *LmexCen^−/−^*-group. This pathway has previously been shown to promote a protective immune response in visceral *leishmania*sis, and could provide a mechanism of protection in the lymph nodes^29^. We also found several other pathways that were uniquely or consistently enriched in the draining lymph nodes of *LmexCen^−/−^*-immunized and *Lmex*WT-infected mice, compared to the naïve group (Supplementary Figure 3). Despite the presence of a few common pathways, the majority of metabolic changes seems to be strain-specific, similarly to what we previously observed in the ears.

## DISCUSSION

American CL is a disfiguring disease that lacks effective preventive strategies. We have recently developed a live attenuated *L. mexicana* parasite lacking the *centrin* gene (*LmexCen^−/−^*) that showed excellent safety and efficacy characteristics in pre-clinical models^12^. In particular, immunization with *LmexCen^−/−^* parasites resulted in enhanced Th1 responses in C57BL/6 mice, contrary to wild type *L. mexicana* infection. In this study, we investigated the metabolic drivers of these immunological profiles.

Although a lot of attention has been given to the role of chemokines and cytokines in *Leishmania* infection, other factors may also influence the immunity against CL. Factors such as microRNAs^30,31^, epigenetic reprogramming, and metabolic mediators^16^ have more recently been investigated. Specifically, metabolomics is emerging as a useful tool to uncover previously unknown signaling networks that govern immune regulation and confer functional specialization. Such metabolic remodeling of immune cell activity has not been extensively explored in vaccine immunity. In particular, live attenuated vaccines harbor significant metabolic differences from their parental strain.

We have previously observed critical differences in the immunological responses following inoculation with *LmexCen^−/−^* parasites compared to their wild type counterparts^12^. This was also observed in *L. donovani, L. major*, and *L. mexicana* centrin mutants^10,12,32,33^. In this study, we explored the metabolic drivers of the immunological differences between infection with the parental or mutant strains and demonstrated that immunization with *LmexCen^−/−^* parasites results in unique metabolic changes, compared to naïve and *Lmex*WT infected mice.

One of the metabolic pathways enriched in the ears of *LmexCen^−/−^* immunized mice, compared to naïve mice, was aspartate metabolism, which is known to induce M1 polarization in macrophages, as well as the production of IL-1β and TNF-α^25^. Consistent with these findings, we observed activated PPP metabolism in the ears of *LmexCen^−/−^* immunized mice, compared to naïve mice, and to mice infected with *Lmex*WT. It is well established that M1 macrophages upregulate glycolysis and PPP in order to generate ATP and NADPH, utilized for the production of ROS and reactive nitrogen species (RNS)^17,20^. Furthermore, the transcription factor Nrf2 has been shown as a regulator of the PPP and a defective PPP has been associated with reduced inflammation^18^. Pro-inflammatory M1 macrophages are crucial for mounting a protective immune response against CL, as they release ROS and RNS (such as NO) to kill the parasites, and act as antigen presenting cells to prime T cells in order to polarize them towards a Th1 phenotype^16,34,35^. The upregulation of aspartate metabolism and PPP at the inoculation site suggests a switch to a protective pro-inflammatory phenotype in macrophages following *LmexCen^−/−^* inoculation. These results are consistent with previous literature showing that an induction in PPP can control the replication of the protozoan parasite *Trypanosoma cruzi* via NO-mediated killing^19^. Interestingly, our results show a PPP-dependent increase in NO, IL-12, and IL-1β production in BMDMs cultured with *LmexCen^−/−^* parasites, which demonstrates a role for PPP in *LmexCen^−/−^*-mediated M1 polarization and pro-inflammatory effector functions in macrophages.

These observations are particularly interesting in light of the fact that *L. mexicana* WT parasites are known to polarize macrophages towards an M2 anti-inflammatory phenotype^15^. This is consistent with previous results showing that *L. donovani Cen^−/−^* parasites also promote an M1 phenotype in macrophages, in contrast to *L. donovani* WT infection^36^.

Furthermore, a recent study showed that DNA release through the process of neutrophil extracellular traps (NETs) was significantly elevated in neutrophils incubated with *L. donovani Cen^−/−^*, compared to *L. donovani* WT parasites^37^, through an unknown mechanism. Interestingly, induction of NETosis requires a shift towards the PPP in neutrophils in order to fuel NADPH oxidase to produce superoxide^38^. Our metabolomics results suggest that the elevated PPP induced by immunization with *centrin^−/−^* parasites could be responsible for elevated NETosis activity in neutrophils, thought that needs to be experimentally demonstrated. Further studies are warranted to determine which cell types predominantly and consistently upregulate aspartate metabolism, as well as the PPP, at the site of inoculation after exposure to *LmexCen^−/−^* mutants. In particular, it would be interesting to determine whether the PPP remains enriched during the adaptive immune response, as this pathway has been shown to be upregulated in activated and proliferating T cells^39^ and it is the signature metabolic pathway of effector T cells^40^.

Our analysis also revealed activated taurine and hypotaurine metabolism in the ears of *LmexCen^−/−^* immunized mice, compared to mice infected with *Lmex*WT. Taurine is known to have anti-apoptotic, antioxidant, and cytoprotective functions^41^, and it promotes wound healing in CL lesions caused by *L. major*^28^. Furthermore, taurine induces the expression of type I IFN-signature genes and promotes phosphorylation of interferon regulatory factor (IRF)-7 in DCs^42^. IRF-7 is crucial for macrophage killing of *L. donovani* in the liver and spleen, and the lack of this factor leads to impaired CD4+ T, natural killer (NK), and NKT cell activity in the liver^43,44^. Additional studies are required to determine the phosphorylation state of IRF-7 and the subsequent immunoregulation following exposure with *LmexCen^−/−^* or *Lmex*WT parasites.

In this study we also analyzed metabolic changes in the draining lymph nodes. Interestingly, the metabolic signatures in the draining lymph nodes at 7 days post infection were somewhat different compared to what we observed at the lesion site. Nevertheless, we found enrichment of linoleic acid (LA) metabolism in *LmexCen^−/−^* immunized mice, compared to mice infected with *Lmex*WT. It has been recently shown that LA inhibits *L. donovani* survival in macrophages and promotes production of superoxide by neutrophils and macrophages^29^. Furthermore, LA is a precursor of ω-6 polyunsaturated fatty acids (PUFAs), and deficiency of PUFAs impairs cell-to-cell interaction, resulting in improper antigen presentation and inadequate lymphocyte activation^29^. Thus, the intake of appropriate amounts of LA through diet can make a difference between resistance and susceptibility to *L. donovani* in patients^29^. Although these results have not been validated in other species of *Leishmania*, they may affect mechanisms of vaccine immunity.

Expectedly, metabolomics analysis indicated several pathways were enriched consistently or uniquely among immunized and infected mice, as illustrated in the Venn diagrams. In this study we focused on the PPP, as well as aspartate, taurine, and linoleic acid metabolism in particular due to their known immunomodulatory functions. In particular, *LmexCen^−/−^*-mediated activation of the PPP lead to the upregulation of pro-inflammatory cytokines in macrophages, indicative of an M1 phenotype. Pro-inflammatory antigen presenting cells play a crucial role in inducing a Th1 adaptive response, which has been previously shown to promote *LmexCen^−/−^*-mediated protection against CL^12^.

In conclusion, to our knowledge this is the first report of a metabolic study done in the context of vaccination against leishmaniasis. Uncovering the unique metabolic signatures of vaccination and disease will be invaluable to identify novel markers of vaccine efficacy against leishmaniasis and other diseases.

## LIMITATIONS

The results in Table 1, show one of the complications of untargeted metabolomics, which is the presence of multiple isoforms of the same compound that can sometimes have opposite fold change directions and create potential ambiguity in the interpretation of the results. Additionally, the annotation of the metabolites (shown with HMDB identifiers) is tentative as they are based exclusively on the formula with no additional information regarding the compound structures.

## MATERIALS AND METHODS

### Mouse strains and parasites

Female mice with a C57BL/6 background were purchased from Envigo (Harlan laboratories) Indianapolis, IN, USA, and housed at The Ohio State University animal facility, following approved animal protocols and University Laboratory Animal Resources (ULAR) regulations (2010A0048-R3 Protocol). All the experiments were performed using 3 to 5 age-matched 5-8 week old female mice per group.

*L. mexicana* (MNYC/B2/62/m379) parasites were maintained via subcutaneous inoculation into the shaved back rumps of 129S6/SvEvTac mice (purchased from Taconic Biosciences, Inc.). Amastigotes extracted from the draining lymph nodes of 129S6/SvEvTac mice were then cultured in complete M199 medium (Gibco, Thermo Fisher Scientific, Waltham, MA), complemented with 1% Penicillin/Streptomycin, 1% HEPES, and 10% fetal bovine serum (FBS) at 26°C to generate stationary phase promastigotes.

### *In vivo* infection

Aged-matched C57BL/6 mice were inoculated intradermally in the ear with 1×10^6^ *L. mexicana* promastigotes in the stationary phase. After 7 days, the ipsilateral infected ear and the naïve ears were collected and processed for mass spectrometry.

### Mass spectrometry

The ear tissue was collected, snap frozen, and processed for mass spectrometry analysis according to SOP 7 of the Laboratory Guide for Metabolomics Experiments^45^. Samples were then incubated with 500 ul of 100% MeOH and sonicated. The tissue was weighed and homogenized at 40 mg/mL of 50% MeOH solution for 3 cycles in a homogenizer with Precellys. The supernatant was collected, dried down, and reconstituted in ½ of the original volume in 5 % MeOH with 0.1% formic acid.

Untargeted analysis was performed on a Thermo Orbitrap LTQ XL with HPLC separation on a Poroshell 120 SB-C18 (2×100 mm, 2.7μm particle size) with an WPS 3000 LC system. The gradient consisted of solvent A, H2O with 0.1% Formic acid, and solvent B 100% acetonitrile at a 200μL/min flow rate with an initial 2 % solvent B with a linear ramp to 95% B at 15 min, holding at 95% B for 1 minutes, and back to 2% B from 16 min and equilibration of 2 % B until min 32. For each sample, 5μL were injected and the top 5 ions were selected for data dependent analysis with a 15 second exclusion window.

For feature selection in the untargeted results analysis, including database comparison and statistical processing, samples were analyzed in Progenesis QI and the pooled sample runs were selected for feature alignment. Anova p-value scores between the groups were calculated with a cutoff of < 0.05. With database matching using the [Human Metabolome Database], selecting for adducts M+H, M+Na, M+K, and M+2H and less than 10ppm mass error, unique features were tentatively identified as potential metabolites.

### Statistical analysis of mass spectrometry datasets

Peak intensity data tables from the mass spectrometry experiment were formatted into comma-separated values (CSV) files conforming to MetaboAnalyst’s requirements and uploaded into the one-factor statistical analysis module. Each analysis passed MetaboAnalyst’s internal data integrity check and additional data filtering was performed based on interquartile range. On the normalization overview page, sample normalization was performed based on the median of the data and the auto-scaling option was chosen to perform data scaling; No transformation of the data was performed. For dimensionality reduction, both principal component analysis (PCA) and partial least-squares discriminant analysis (PLS-DA) were employed. Cross-validated sum of squares (Q2) performance measures were used to determine if PLS-DA models were overfitted. Visualization of significant, differentially regulated metabolites was done by generating volcano plots with cutoffs of <0.05 false-discovery rate (FDR) and >2-fold change (FC). Clustering of samples and features were analyzed by creating dendrograms and hierarchical heatmaps, respectively.

### Pathway analysis of mass spectrometry datasets

We have used two different techniques in order to identify enriched pathways in our data sets. First, we used the Functional Analysis Module (MS peaks to pathways) in MetaboAnalyst 4.0. Detected peaks (mass-to-charge ratios + retention times) from positive and negative analytical modes of the mass spectrometer for each sample were organized into four column lists along with calculated p-values and t-scores from univariate t-tests. These peak list profiles were uploaded to the functional analysis module and passed the internal data integrity checks. The ion mode in MetaboAnalyst was set to the appropriate type depending on the analytical mode that was used to generate the data. For each analysis the mass tolerance was set to 10 ppm, the retention time units were set to minutes, and the option to enforce primary ions was checked. In parameter settings, the mummichog algorithm (version 2.0) and the modified gene set enrichment algorithm were used for all analyses. The p-value cutoff for the mummichog algorithm was left at the default (top 10% of peaks). Currency metabolites and adducts were left at default settings. Lastly, the Kyoto Encyclopedia of Genes and Genomes (KEGG) pathway library for *Mus musculus* was selected as the metabolic network that the functional analysis module would use to infer pathway activity and predict metabolite identity; only pathways/metabolite sets with at least three entries were allowed.

To confirm the enriched pathways identified with MetaboAnalyst, we also used the Ingenuity Pathway Analysis (IPA) software. Metabolite matches to the detected peaks from database searches (HMDB and LIPID MAPS), along with calculated p-values and fold changes were uploaded into IPA software for core analysis. The reference set was selected as the Ingenuity Knowledge Base (endogenous chemicals only) and direct and indirect relationships were considered during analysis. Settings in the networks, node type, data sources, confidence, and mutations tabs were left at default values. The species tab settings were set to mouse and uncategorized (selecting uncategorized species for metabolomics is necessary in IPA as most metabolites are not unique to any one species). Lastly, in the tissues and cell lines tab all tissues were considered in the analysis, whereas all cell lines were excluded from consideration. The p-value cutoff for every analysis was set as 0.05 and the fold change cutoffs were adjusted to obtain between 200-1000 analysis-ready molecules and then kept the same across different analyses.

### *In vitro* cell culture and infection

Bone marrow-derived macrophages (BMDMs) were obtained from the femur and tibias of C57BL/6 mice. After isolation, the bone marrow cell suspension was cultured with RPMI medium supplemented with 10% fetal bovine serum (FBS), 1% penicillin/streptomycin, 1% HEPES, and 20% supernatant from L-929 cells at 37°C with 5% CO_2_ for 7-10 days until differentiation was complete. BMDMs were then plated in a 24-well plate at a density of 0.5×10^6^ per well and infected overnight with stationary phase *LmexCen^−/−^* promastigotes at a ratio of 10:1 (parasite to macrophages). The naïve controls were treated with media alone. Then, the extracellular parasites were removed by washing with PBS and new media was applied. Some of the groups were stimulated with LPS (1μg/ml) for 24 hrs after infection and then washed with PBS. Some of the groups were then treated with 100μM 6-aminonicotinamide (6-AN) or dehydroepiandrosterone (DHEA) (both from Sigma-Aldrich), two pentose phosphate pathway inhibitors^18^. After a 24hrs incubation, the supernatant was collected to measure nitric oxide and cytokine levels.

### Nitric oxide assay

Nitric oxide release from BMDMs was determined in the culture supernatants by measurement of nitrite using Griess reagent (Sigma Aldrich) and sodium nitrite as the standard. A Molecular Devices SpectraMax M3 microplate reader was used to measure the absorbance at 570 nm, and concentrations of nitric oxide were determined from the standard curve with the Softmax Pro software (Molecular Devices LLC).

### Cytokine ELISA

Levels of IL-12 and IL-1β in BMDM supernatants were determined by sandwich ELISA. 96-well plates were coated with primary capture antibody against IL-12 (Biolegend Cat#511802, clone C18.2) and IL-1β (Biolegend Cat#503502, clone B122) at a final concentration of 2μg/ml, and incubated overnight at 4°C. Plates were then blocked with PBS + 10% FBS for 2 hrs at RT and incubated overnight with 50 μl of culture supernatants or recombinant cytokine (standard curve) in duplicates at 4°C. After washing with PBS-Tween (0.05% Tween 20 in 1× PBS, pH 7.4), the plates were incubated with biotinylated detection antibodies against IL-12 (Biolegend Cat#505302, clone C17.8) and IL-1β (Biolegend Cat#515801, clone Poly5158) at a final concentration of 1μg/ml, for 1 hr at RT, washed again, and incubated with streptavidin-conjugated Alkaline Phosphatase for 30 mins at RT in the dark. After washing again, the plates were incubated with PNPP until development. A Molecular Devices SpectraMax M3 microplate reader was used to measure absorbance at 405 nm, and the SoftMax Pro software was used to quantify cytokine levels against the standard curve. All reagents for ELISA were purchased from Biolegend Inc.

### Statistical analysis

All *in vitro* and *in vivo* data show a representative experiment with N ≥ 3 per group. N represents different biological replicates. For the mass spectrometry data all statistical analysis were performed with MetaboAnalyst and IPA software. For the nitric oxide assay, unpaired two-tailed Student’s t test was performed to compare statistical significance. A p value < 0.05 was considered significant. In all figures * represents p ≤ 0.05, ** represents p ≤ 0.01 and *** represents P ≤ 0.001. Error bars represent SEM (standard error of the mean).

## DATA AVAILABILITY STATEMENT

All relevant data is available in the main text and supplementary information. Any additional information can be provided upon reasonable request to the authors.

## AUTHOR CONTRIBUTIONS

G.V., S.G., H.L.N., and A.R.S. designed the experiments. G.V. performed the experiments. T.O. and N.A. analyzed the mass spectrometry data. G.V. and S.G. wrote the manuscript. G.V., T.O., N.A., S.G, H.L.N, S.H., G.M., and A.R.S. revised the manuscript.

## FIGURES AND LEGENDS

**Supplementary Figure 1.**
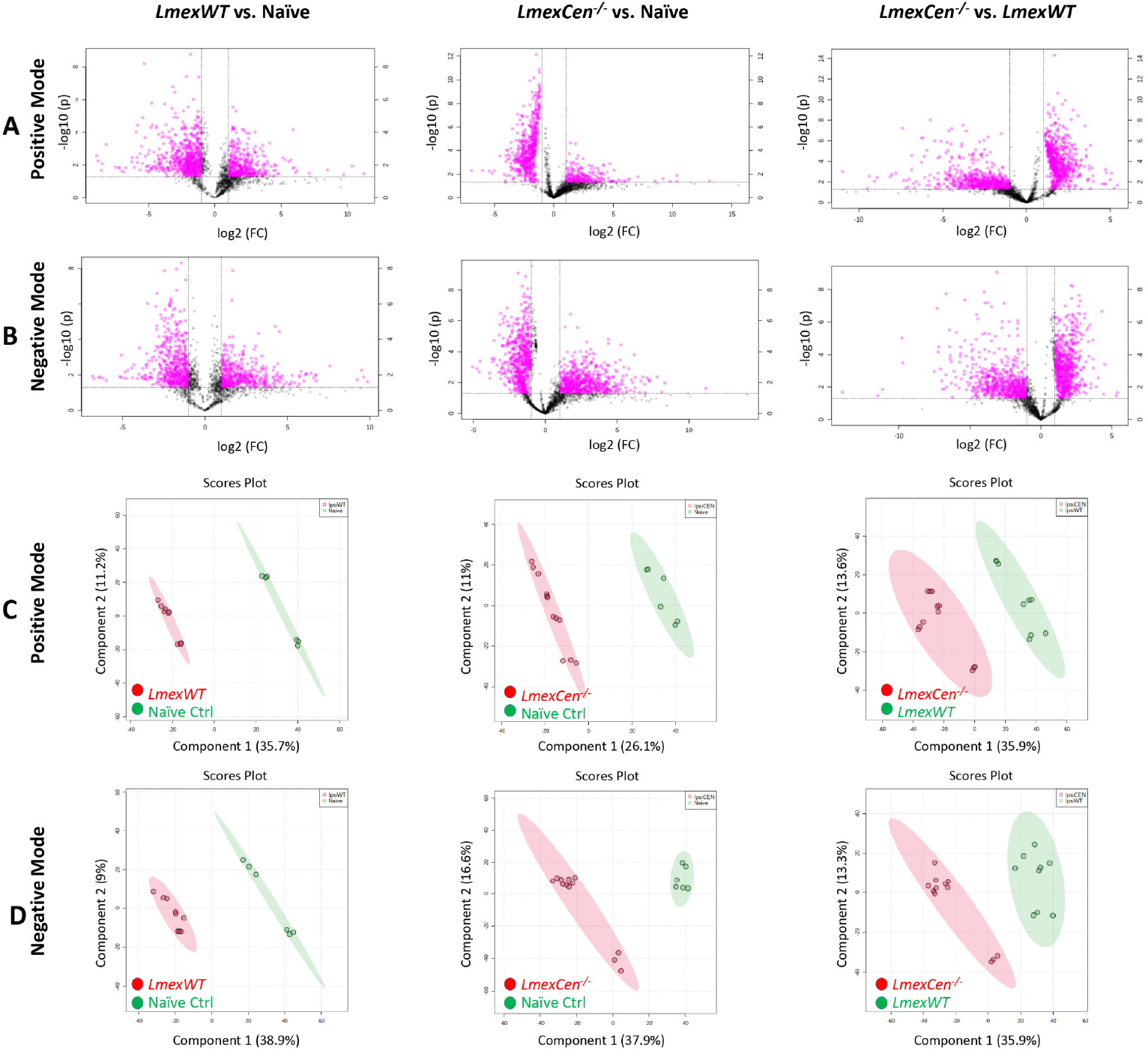
Infection with *Lmex*WT and immunization with *LmexCen^−/−^* display different metabolic signatures in the draining lymph nodes. Normalized data from draining lymph nodes from C57BL/6 mice after 7 days of infection with *Lmex*WT, inoculation with *LmexCen^−/−^*, or naïve control was used to perform statistical analysis. **A, B)** Features selected by volcano plot from positive (**A**) and negative (**B**) modes for draining lymph nodes of *Lmex*WT vs. naïve, *LmexCen-/-* vs. naïve, and *LmexCen-/-* vs. *Lmex*WT mice using LC/MS with fold change threshold (x) 2 and t-tests threshold (y) 0.05. Both fold changes and p-values are log transformed. **C, D)** PLS-DA from positive (**C**) and negative (**D**) mode for draining lymph nodes of *Lmex*WT vs. naïve, *LmexCen-/-* vs. naïve, and *LmexCen-/-* vs. *Lmex*WT mice.

**Supplementary Figure 2.**
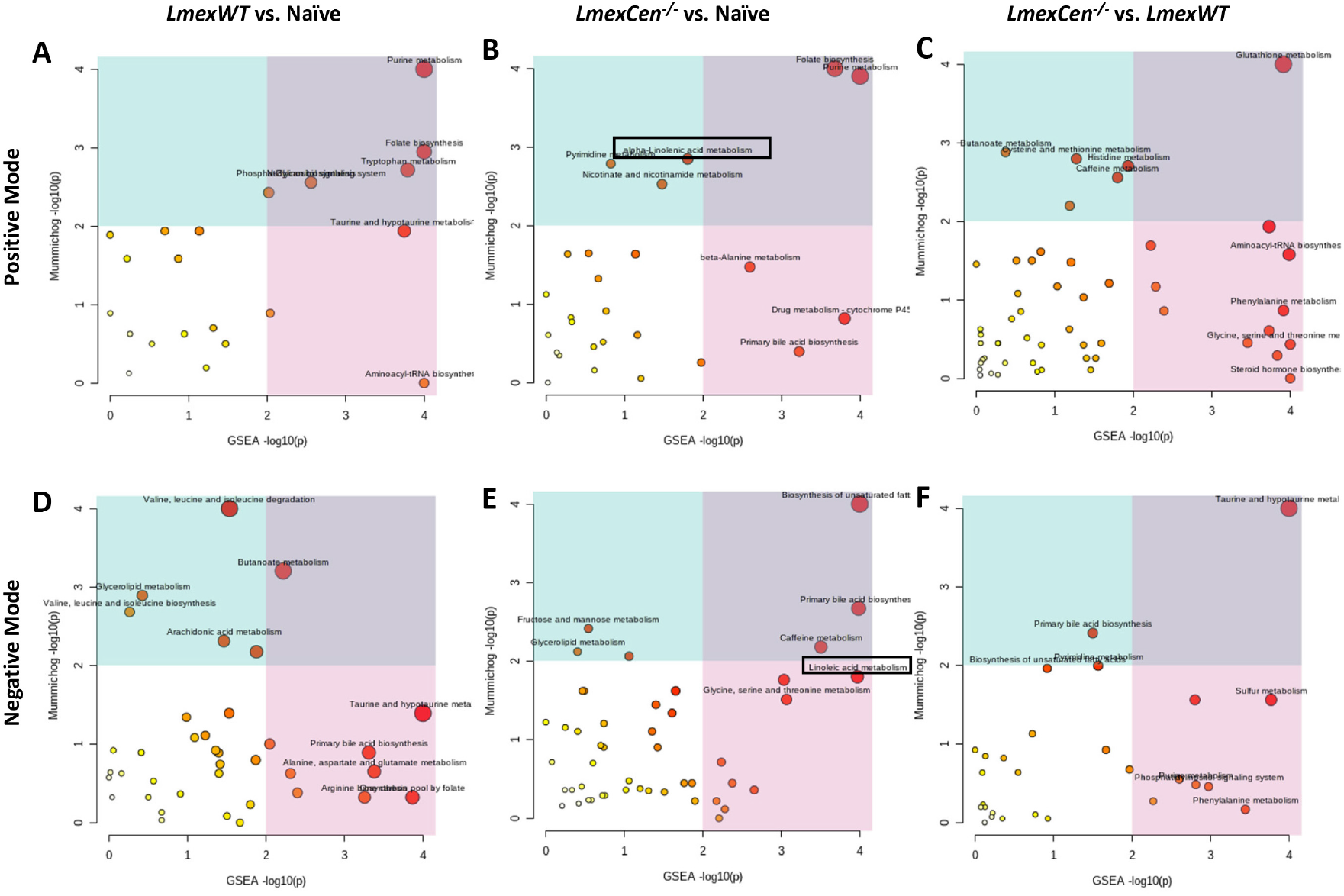
Metabolic pathways enriched in mice infected with *Lmex*WT or immunized with *LmexCen^−/−^* in the draining lymph nodes. Normalized data from draining lymph nodes from C57BL/6 mice after 7 days of infection with *Lmex*WT, immunization with *LmexCen^−/−^*, or naïve control was used to perform peaks to pathway analysis. Using MS Peaks to Paths module in MetaboAnalyst5.0, the mummichog and GSEA p-values are combined. The Integrated MS Peaks to Paths plot summarizes the results of the Fisher’s method for combining mummichog (y) and GSEA (x) p-values from the positive (**A**) negative (**B**) mode data sets, indicating the metabolic pathways enriched. The size and color of the circles correspond to their transformed combined p-values. Large and red circles are considered the most perturbed pathways. The colored areas show the significant pathways based on either mummichog (blue) or GSEA (pink) and the purple area highlights significant pathways identified by both algorithms. Highlighted by the black boxes are pathways associated with a protective immune response against leishmaniasis.

**Supplementary Figure 3.**
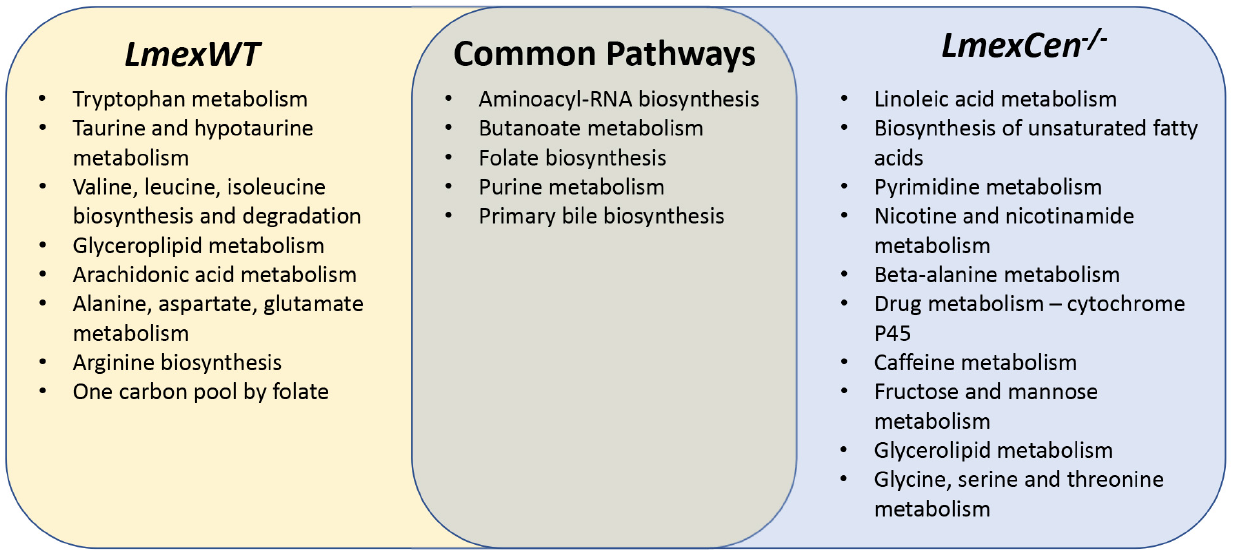
Unique and common pathways enriched in the draining lymph nodes following *LmexCen^−/−^* immunization or *Lmex*WT infection. Venn diagrams of the metabolic pathways enriched based on mummichog and GSEA analysis in the draining lymph nodes of C57BL/6 mice after 7 days of infection with *Lmex*WT or immunization with *LmexCen-/-*, compared to uninfected naïve mice.

